# A novel transformer-based platform for the prediction and design of biosynthetic gene clusters for (un)natural products

**DOI:** 10.1101/2025.06.02.657346

**Authors:** Tomoki Kawano, Taro Shiraishi, Tomohisa Kuzuyama, Maiko Umemura

## Abstract

Biosynthetic gene clusters (BGCs), comprising sets of functionally related genes responsible for synthesizing complex natural products, are a rich source of bioactive compounds with pharmaceutical potential. Here, we present a transformer-based framework that models functional domains as linguistic units to capture and predict their positional relationships within genomes. Using a RoBERTa architecture, we trained models on four progressively broader datasets: bacterial BGCs, Actinomycetes genomes, bacterial genomes, and bacterial plus fungal genomes. Evaluation using 2,492 experimentally-validated BGCs from the MIBiG database showed that more than 60% of true domains were ranked first and over 80% within the top 10 candidates. Our models also achieved classification accuracies exceeding 70% for major compound classes including polyketides (PKs) and terpenes. To explore model-guided BGC design, we compared predictions from the BGC-trained and genome-trained models using the BGC for the bacterial diterpenoid cyclooctatin as a case study. The genome-trained model uniquely predicted several domains absent from both the original BGC and the prediction by the BGC-trained model. Heterologous expression of one of those predicted domains in *Streptomyces albus*, together with the biosynthetic genes for cyclooctatin, yielded an unknown cyclooctatin derivative. This framework not only provides a novel BGC prediction method using machine learning but also facilitates rational design of artificial BGCs. Future integration of transcriptomic, protein structural, and phylogenetic data will enhance the models’ predictive and generative capabilities, supporting accelerated discovery and engineering of natural products.

**Author Summary:** BGCs encode diverse natural products, including antibiotics and anticancer agents. Identifying and designing BGCs in microbial genomes is crucial for discovering new bioactive compounds. In this study, we developed a transformer-based deep learning model that treats protein domains as language-like tokens and learns how they are arranged in genomes. By training on both known BGCs and whole genomes, the model successfully predicts biologically plausible combinations of domains, including those absent in known BGCs. We experimentally validated one such prediction by expressing a newly identified gene alongside known cyclooctatin biosynthetic genes, confirming the production of an unknown cyclooctatin derivative. Our results demonstrate how language models can uncover hidden biosynthetic potential and offer a promising new AI tool for natural product discovery and synthetic biology.

## Introduction

Natural products have provided many important drugs such as penicillin, cyclosporine, tacrolimus, and paclitaxel, making them invaluable pharmaceutical resources. These compounds, primarily produced by microorganisms and plants, possess complex chemical structures that are often challenging to synthesize using conventional methods. The genes responsible for producing these compounds typically co-localize in biosynthetic gene clusters (BGCs), representing genomic regions where functionally related genes cooperate to produce specific natural products.

The organization of BGCs reflects their evolutionary history. In prokaryotes, multiple genes in a BGC often form operons regulated by one or a few promoters. In contrast, eukaryotes lack operon structures, but BGC gene expression is frequently coordinated by transcription factors encoded within the cluster. Recent experimental work by Kanai et al. [1] showed that DNA rearrangement via transposases facilitates operon formation, providing insights into BGC evolution. These evolutionary events leave genomic footprints as differences in the gene organization within cluster structures, as seen in cyanobactin BGCs across *Spirulina* species (Fig 1). Cyanobactins are cyclic peptides produced by cyanobacteria, some of which exhibit antitumor activity [2]. In the figure, the top BGC contains a transposase gene and four endonuclease genes, whereas the bottom three lack these elements. The second and third BGCs retain endonucleases but lack transposases. Additionally, gene orientation shifts from random to uniform directionality. These patterns imply that the evolutionary history of BGC formation is embedded in their genomic context, analogous to a recording device.

**Fig 1.**
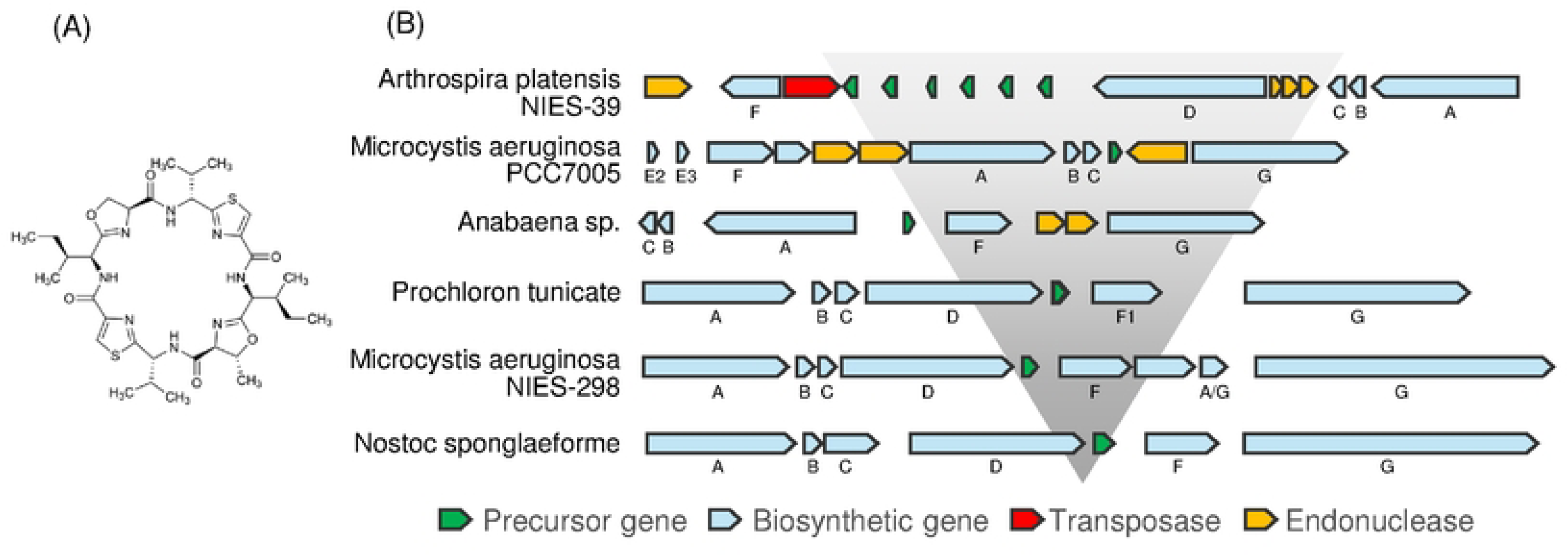
Gene organization in cyanobactin biosynthetic gene clusters (BGCs) from cyanobacterial strains. (A) Structure of patellamide A, a representative cyanobactin produced by *Prochloron didemni*. (B) Comparison of cyanobactin BGCs from six different cyanobacterial strains, showing variation in gene content and arrangement.

Recent advances in genome sequencing and computational methods have significantly improved BGC identification [3]. Current computational tools like antiSMASH [4] use hidden Markov models to detect BGCs through functional genes specific to natural compound biosynthesis, but they largely depend on known domain patterns and are limited in their ability to discover novel BGC architectures. Deep learning-based tools such as DeepBGC [5] have emerged, yet they also rely heavily on pre-defined BGC features.

Advances in artificial intelligence, particularly in natural language processing, provide new opportunities for analyzing genomic sequences. Transformer-based models that are superior at learning long-range dependencies in text have proven useful in biology, especially in protein structure prediction and functional annotation [6-8]. Large-scale models such as Evo have demonstrated the ability to predict and generate genomic sequences at megabase scales [9]. These examples suggest that transformer models may be well-suited for capturing the positional and contextual relationships among functional domains in genomes.

In this study, we introduce a transformer-based framework for predicting and designing BGCs by learning the sequential organization of functional domains as a proxy for evolutionary constraints. Functional domains are treated as language-like tokens, and domain arrangements are modeled using natural language processing techniques. A distinguishing feature of our approach is that it learns from both BGC and non-BGC genomic contexts, enabling the discovery of novel domain combinations that are difficult to identify using existing tools. Our models, currently trained solely on static representations of functional domain sequences in genomes, will later be fine-tuned using multimodal information related to evolutionary direction such as transcriptomic data and compound productivity, aiming to generate more evolutionarily informed or functionally optimized BGCs for specific biosynthetic objectives.

## Results

### Pretraining on Genomes Using Functional Domain-Based Tokenization

We adopted functional domains within genes as token units for modeling genomic sequences, based on their established importance in BGC characterization as utilized in bioinformatics tools such as antiSMASH. We employed the RoBERTa architecture [10] for this purpose, configured with 8 hidden layers and 16 attention heads, and a maximum token length of 512 to cover the context of entire BGC domain sequences (Fig 2). We chose the BERT-based architecture due to its bidirectional attention mechanism, which captures context in both directions from each token [11,12]. As gene order in a BGC is often not essential for compound biosynthesis, GPT-style models with unidirectional attention are less suitable for BGC modeling. As shown in Fig S1, both the RoBERTa and GPT-1 models demonstrated effective convergence on Dataset II (described in the following paragraph). However, the GPT-1 model exhibited earlier saturation of the validation loss and failed to produce meaningful predictions for domain tokens in BGC sequences. These results indicate that the bidirectional attention mechanism in RoBERTa offers an advantage in learning the genomic arrangement of functional domains.

**Fig 2.**
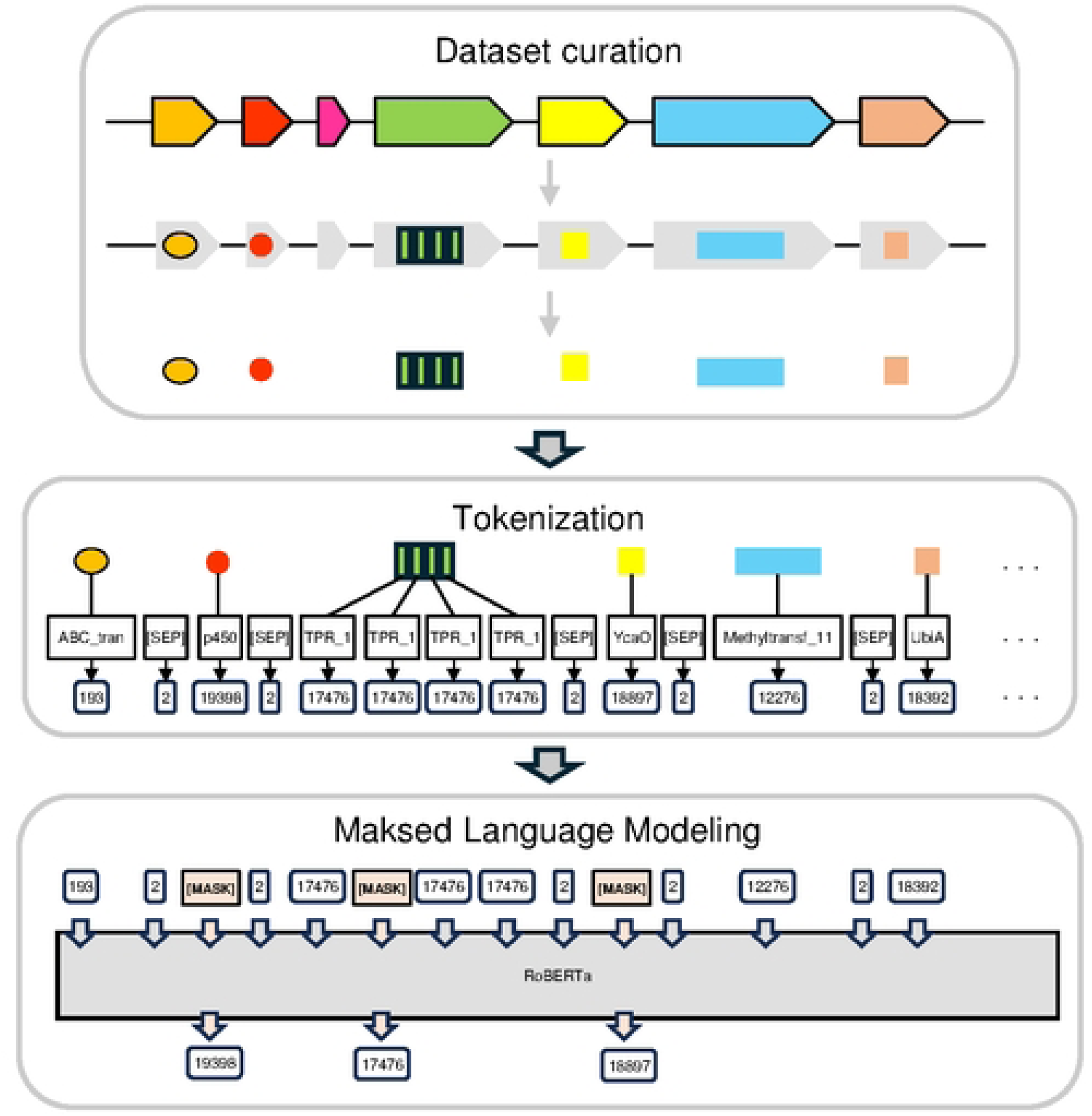
Workflow of masked language modeling of genomic data using RoBERTa. The workflow illustrates the process of tokenizing genomic data and training the RoBERTa language model. In the data curation phase, genomic sequences are retrieved from NCBI and analyzed using HMMer to identify Pfam domains. The tokenization phase converts all identified Pfam domains into discrete tokens. During the masked language modeling phase, these tokenized sequences serve as input for RoBERTa model training, where the model learns to predict masked tokens based on their surrounding context information.

To investigate the impact of training data on model performance, we prepared four datasets of increasing taxonomic coverage: (I) bacterial BGCs from the antiSMASH database; (II) complete genomes of Actinomycetes; (III) complete bacterial genomes; and (IV) complete bacterial plus fungal genomes (Table 1). This stepwise expansion (Fig 3) allowed us to explore how different genomic contexts influence model generalization while maintaining the domain-based tokenization approach consistent.

**Fig 3.**
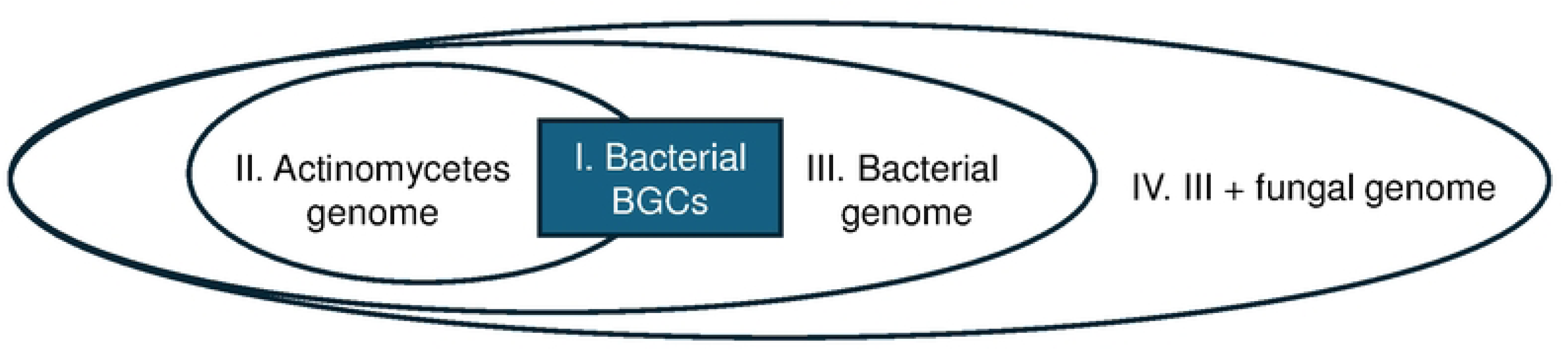
The relationships among four datasets used in this study. The outermost dataset (IV) comprises bacterial and fungal genomes (70.5M tokens from 11,884 strains), encompassing the bacterial genome dataset (III; 48.9M tokens, 9,748 strains) and the Actinomycetes dataset (II; 13.8M tokens, 2,664 strains). The BGC dataset from antiSMASH database (I; 8.6M tokens from 239,021 BGCs) is positioned centrally, reflecting its specialized nature in secondary metabolite biosynthesis.

**Table 1.**
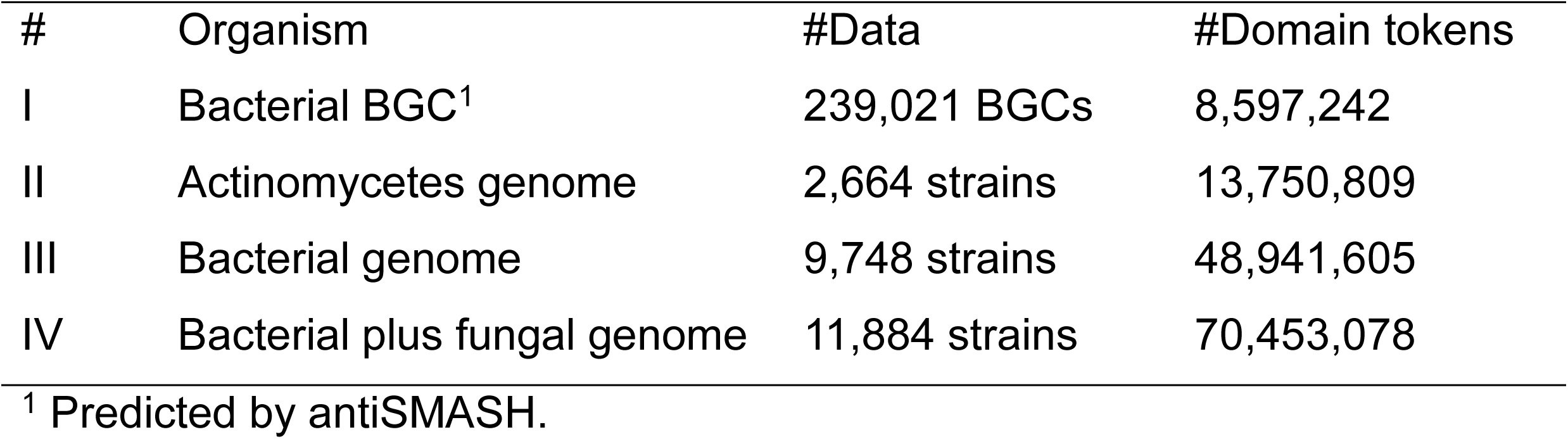
Datasets used for training transformer-based models.

Training on all four datasets resulted in a consistent decrease in validation loss, indicating that the model learns the genomic arrangement of functional domains across the different dataset (Fig 4). The BGC-only model exhibited the most rapid convergence, likely due to its narrower functional diversity. In contrast, the model trained on bacterial plus fungal genomes exhibited the highest validation loss, reflecting the increased complexity of mixed-kingdom data. These results indicate that data complexity directly influences convergence behavior of validation loss during model training.

**Fig 4.**
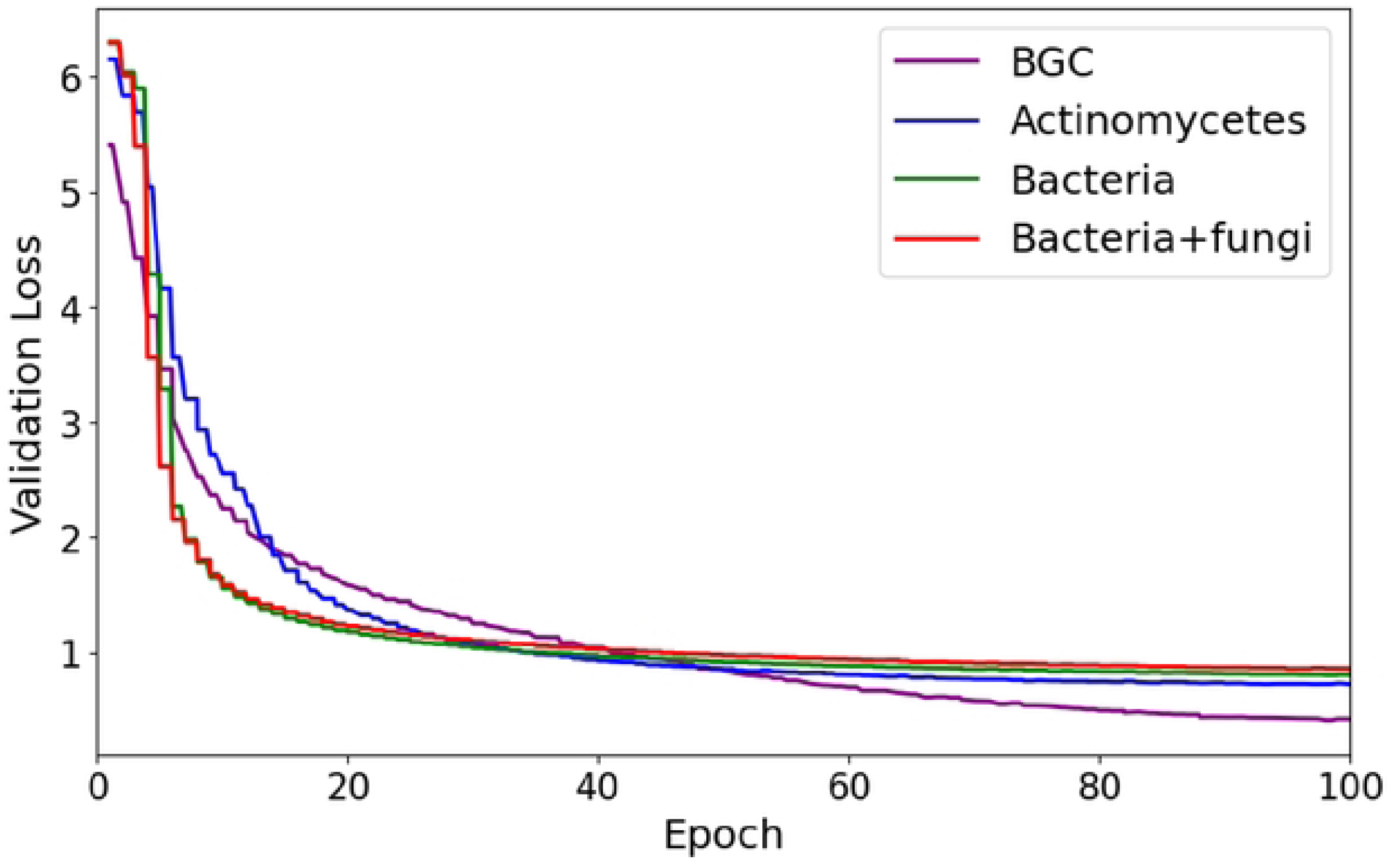
Comparison of validation loss across datasets during model training. Validation loss is plotted over training epochs for four datasets: Biosynthetic gene cluster (BGC), Actinomycetes genome, bacterial genome, and bacterial plus fungal genome. The BGC dataset exhibited the largest reduction in validation loss, likely reflecting its simpler and more homogeneous data structure compared to genomic data. Conversely, the bacterial plus fungal genome dataset exhibited the highest validation loss, likely due to its greater complexity arising from the diverse biological species it encompasses. These results suggest that dataset complexity directly influences the convergence behavior of validation loss during model training.

### Model Performance in BGC Classification

To evaluate the classification performance of the constructed models across different BGC types, we conducted a compound class prediction task using compound class labels from the antiSMASH database (Fig 5). Our model achieved more than 70% classification accuracy for approximately half of the compound classes, with highest performance observed in well-characterized classes such as the Type II polyketide synthase class.

**Fig 5.**
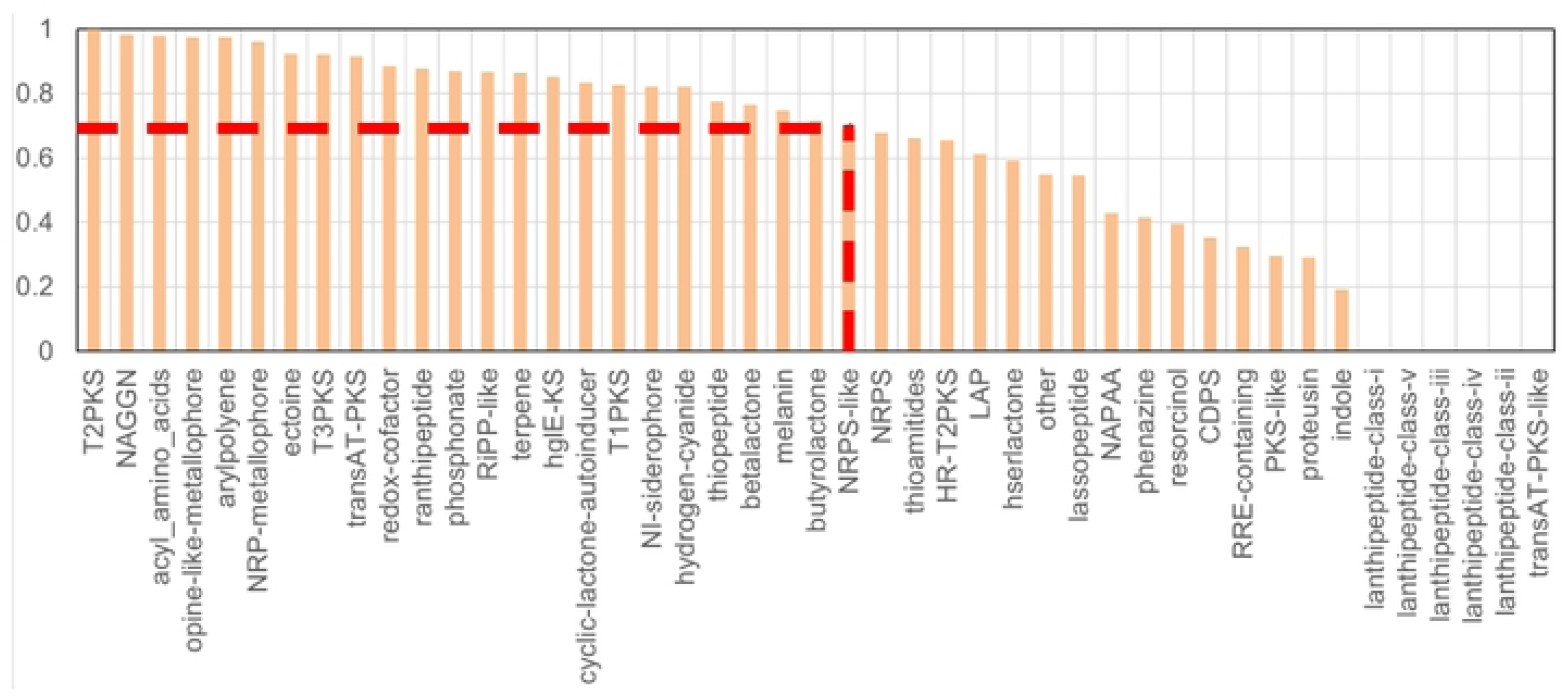
Classification accuracy across compound classes in the antiSMASH database. Bars represent the prediction accuracy of the model for various biosynthetic gene cluster (BGC) classes. The model, trained on 22,258 Actinomycetes BGCs in the antiSMASH database, was evaluated on an independent test set of 5,600 Actinomycetes BGCs. Highest accuracy was observed for classes such as T2PKS and NRPS-like (>0.8). The red dashed line denotes 0.7 accuracy threshold, above which more than half of the compound classes were successfully predicted.

To visualize the model’s capability to distinguish between BGC types, we projected learned feature representations of 2,492 experimentally validated BGCs from the MIBiG database [13] using t-distributed stochastic neighbor embedding (t-SNE) dimensionality reduction [14] (Fig 6). In Fig 6A, major compound classes such as NRP (red) and ribosomally synthesized and post-translationally modified peptide (RiPP, brown) form clearly separated clusters, reflecting the model’s ability to capture class-specific features. In Fig 6B, BGCs associated with polyketide synthases (PKSs) further resolved into Type I (blue) and Type II (red) subclasses. These results support the model’s capability to capture functional and structural characteristics relevant to compound class classification.

**Fig 6.**
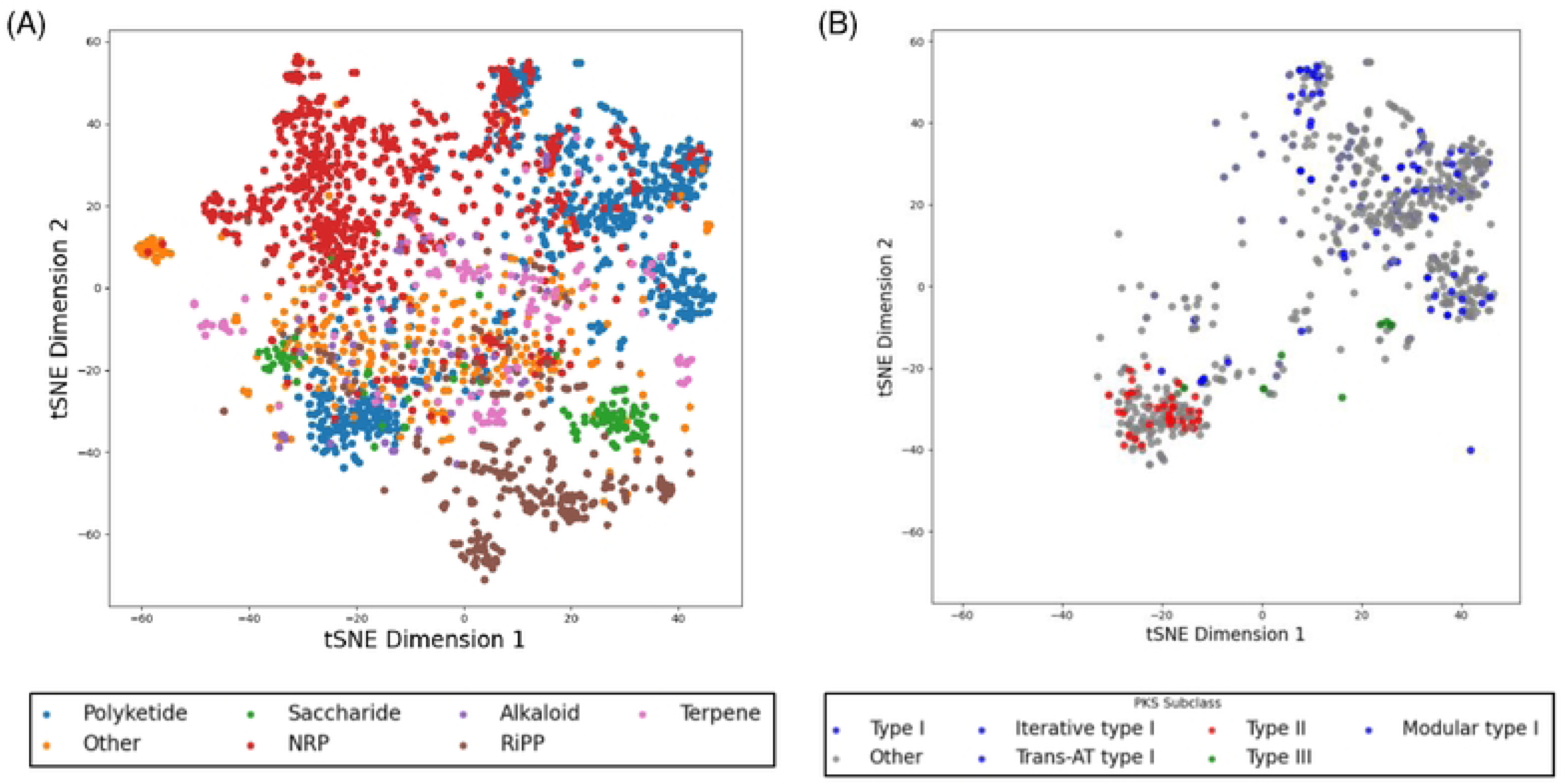
t-SNE visualization of biosynthetic gene cluster (BGC) feature representations. Feature embeddings of BGCs from the MIBIG database were visualized using t-distributed stochastic neighbor embedding (t-SNE). (A) Visualization of all 2,492 BGCs, classified into seven compound classes. Nonribosomal peptides (NRPs, red) and ribosomally synthesized and post-translationally modified peptides (RiPPs, brown) formed distinct clusters. (B) Visualization of polyketide BGCs by subclass. A clear separation is observed, particularly between Type I and Type II polyketide synthases (PKSs).

### Model Performance in Functional Domain Prediction

To evaluate how well the developed models understand domain context, we conducted a masked prediction task using 2,492 BGCs from the MIBiG database. One domain at a time was masked, and models were evaluated on their ability to rank the true domain among the prediction.

Across all models, more than 60% of true domains were top-ranked, and over 80% appeared within the top 10 predictions (Table 2). As training data expanded from Actinomycetes (Model II) to all bacteria (Model III) and then to bacteria plus fungi (Model IV), the average and standard deviation of the prediction ranks improved (Fig 7). These results indicate that the models accurately capture BGC context and positional relationships among functional domains.

**Fig 7.**
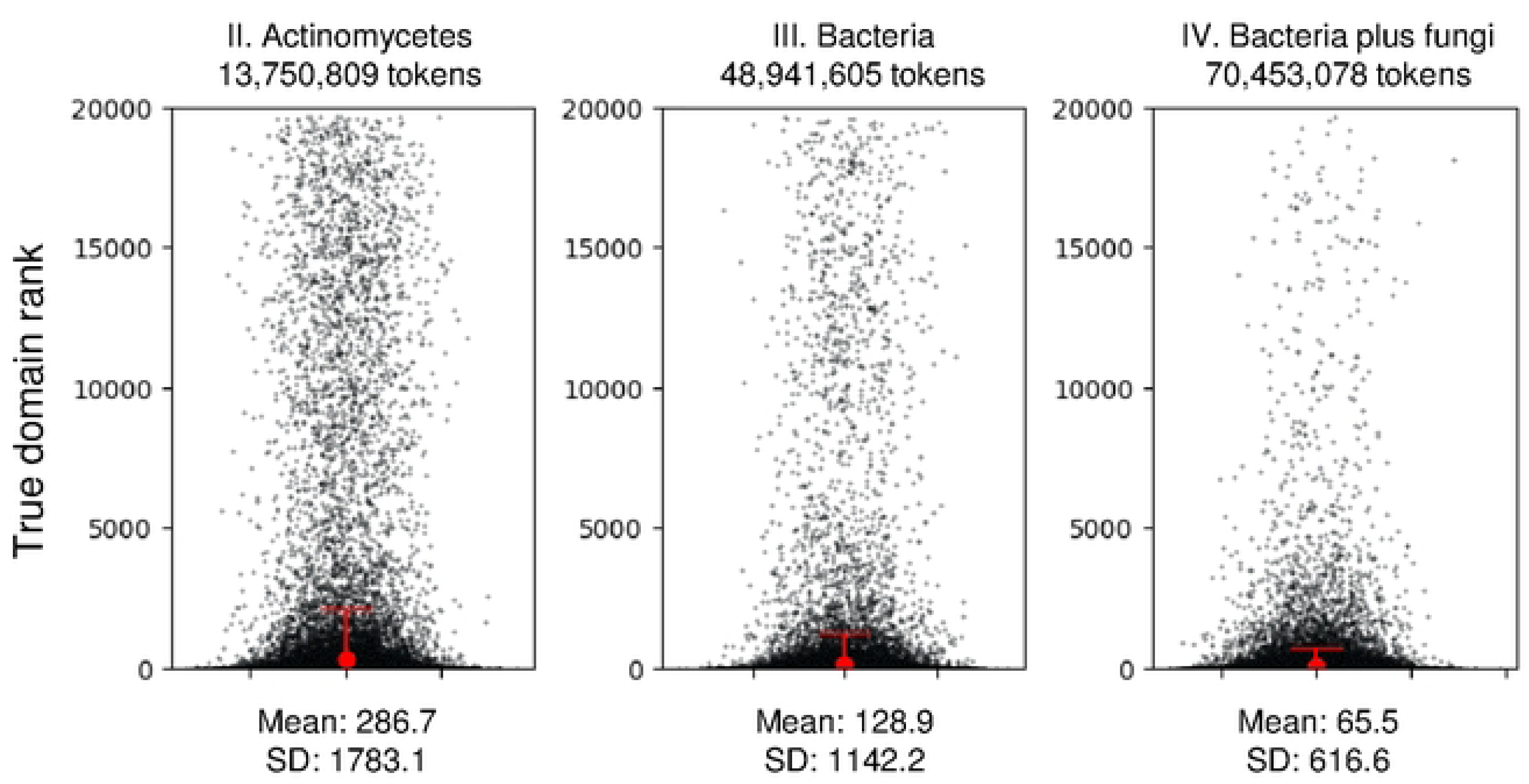
Distribution of prediction rankings for biosynthetic gene clusters (BGCs) from the MIBiG database across four training datasets. Shown are the distributions of true functional domain ranks predicted by models trained on actinomycete genomes (Model II), bacterial genomes (Model III), and bacterial plus fungal genomes (Model IV). Red dots represent the average prediction rank, and red bars indicate the standard deviation (SD) for each dataset.

**Table 2.**
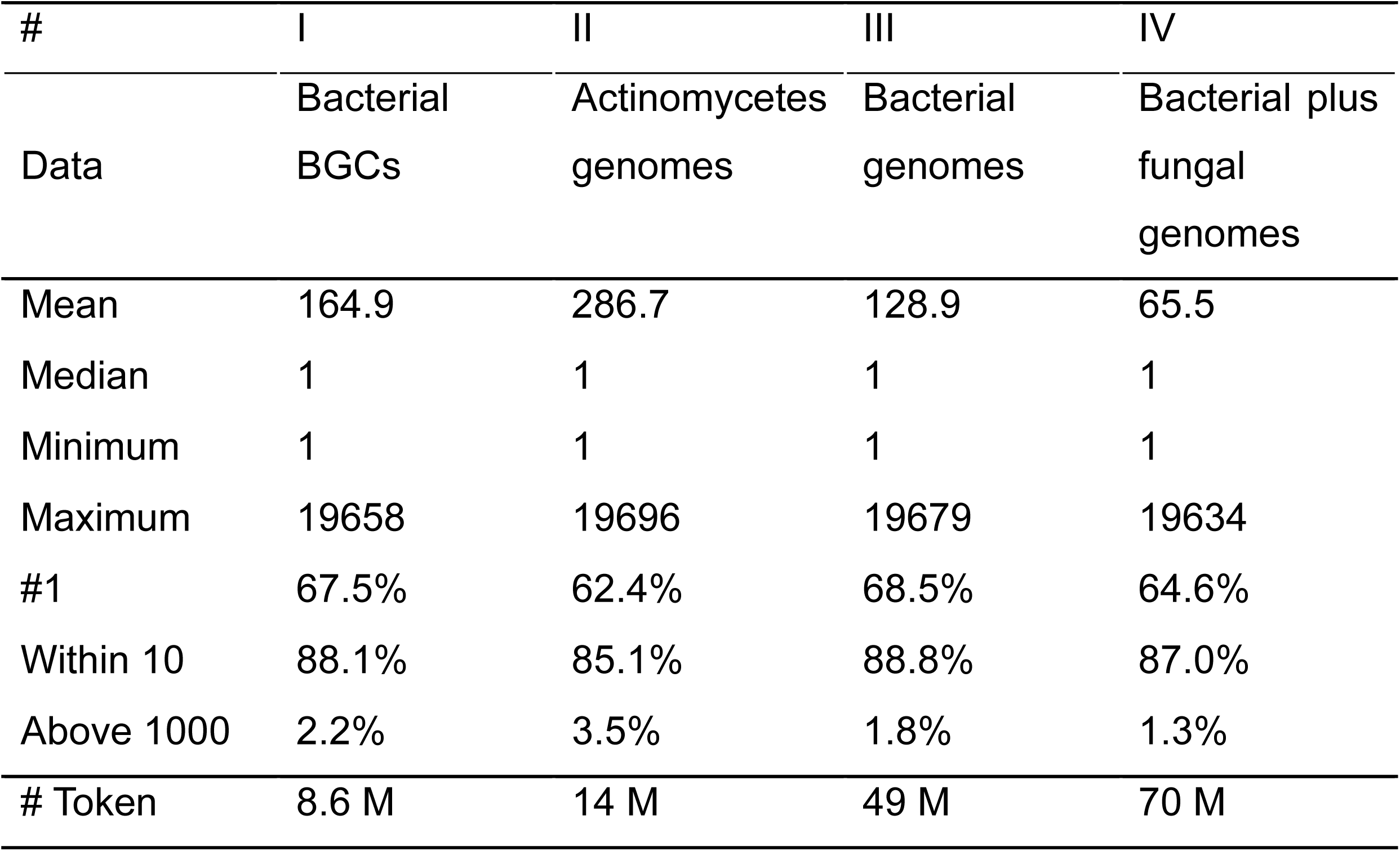
Domain prediction performance across four training datasets using 2,492 biosynthetic gene clusters (BGCs) in the MIBiG database.

| # | I | II | III | IV |
| --- | --- | --- | --- | --- |
| Data | Bacterial BGCs | Actinomycetes genomes | Bacterial genomes | Bacterial plus fungal genomes |
| Mean | 164.9 | 286.7 | 128.9 | 65.5 |
| Median | 1 | 1 | 1 | 1 |
| Minimum | 1 | 1 | 1 | 1 |
| Maximum | 19658 | 19696 | 19679 | 19634 |
| #1 | 67.5% | 62.4% | 68.5% | 64.6% |
| Within 10 | 88.1% | 85.1% | 88.8% | 87.0% |
| Above 1000 | 2.2% | 3.5% | 1.8% | 1.3% |
| # Token | 8.6 M | 14 M | 49 M | 70 M |

However, we observed a slight decrease in top-1 accuracy from 68.5% to 64.6% when fungal data were included (Table 2), possibly due to taxonomic interference in predicting bacterial BGCs. Indeed, when the four models were tested using only the Actinomycetota 1,042 BGCs from MIBiG, both top-1 accuracy and the mean prediction rank declined: the mean rank increased from 31.8 to 35.4 upon inclusion of fungal data (Table S1).

### Context Dependency of Domain Prediction

To determine how input context affects domain prediction by the constructed models, we performed a systematic perturbation analysis in which input domains were randomly altered. We observed the prediction probability of the true domain at a masked position domain by Model IV as increasing the number of altered domains (Fig 8).

**Fig 8.**
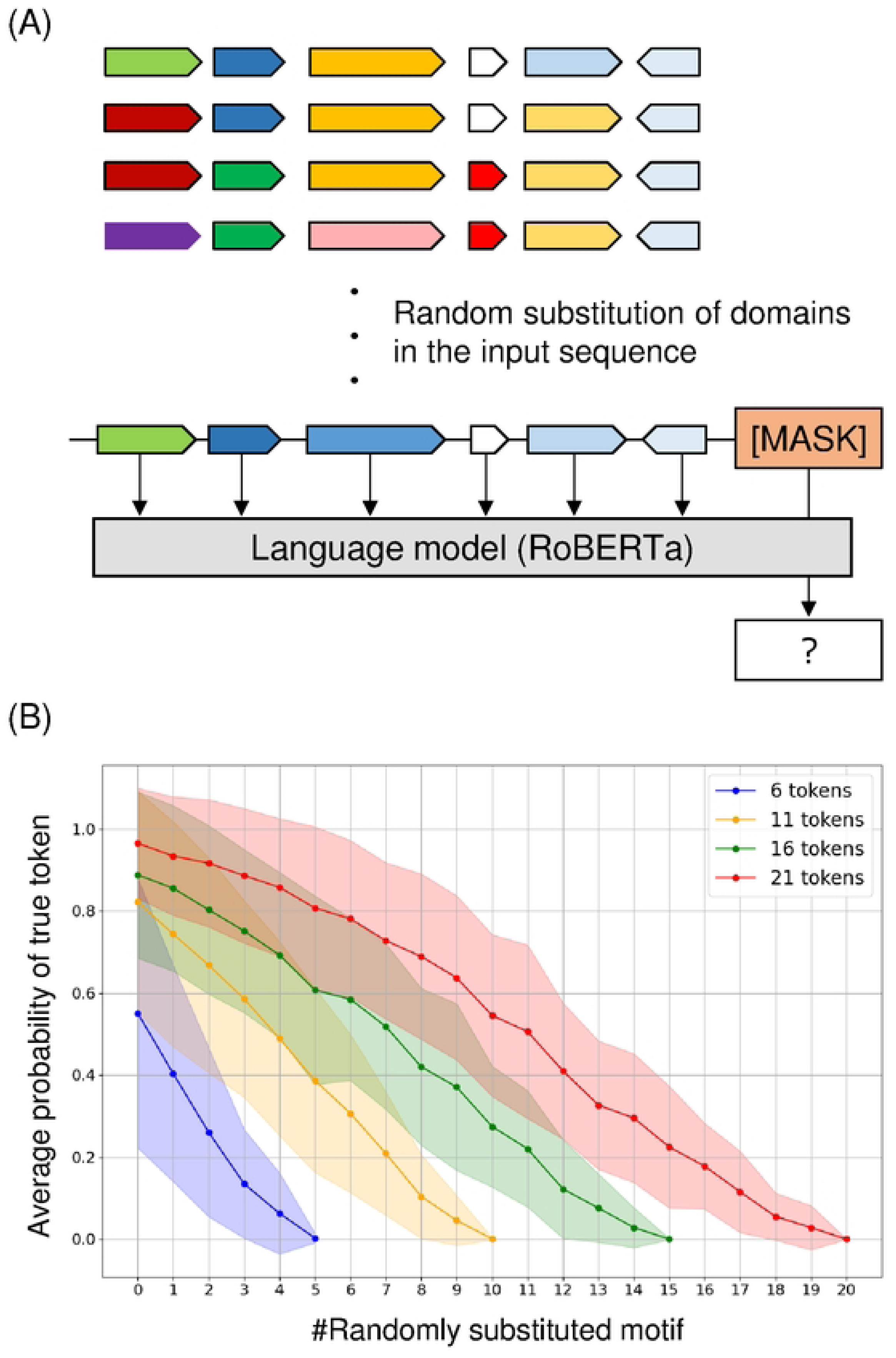
Effect of token substitutions on domain prediction accuracy. (A) Schematic of the perturbation experiment to investigate how prediction accuracy for a masked token changes when randomly substituting other tokens in the input sequence (excluding [SEP] tokens). Colored arrows represent functional domains in proteins. (B) Prediction results for domain clusters consisting of 6, 11, 16, and 21 tokens (sample sizes of 81, 66, 71, and 65 clusters, respectively). The y-axis shows the model’s prediction probability for the correct token at the [MASK] position, while the x-axis indicates the number of randomly substituted tokens. Shaded areas represent standard deviations across samples.

The average of prediction probability decreased monotonically as replacement rate increased, across BGCs containing 6, 11, 16, and 21 functional domains from the MIBiG database (Fig 8B). These results indicate that unmasked domains serve as critical contextual cues, effectively guiding the model in predicting masked BGC domains.

### Design of Artificial BGCs from Model Prediction

We next tested the model’s generative capability to propose plausible domain combinations. Compared to the model trained solely on known BGCs, the model trained on comprehensive genome datasets is expected to predict broader and more diverse sets of biologically plausible domain combinations, including those not observed in known BGCs. To illustrate this, we compared the candidate lists generated by Model I (trained only on BGCs) and Model IV (trained on bacterial plus fungal genomes) using the cyclooctatin biosynthetic pathway as a case study. Cyclooctatin is a C_20_ diterpene produced by *Streptomyces melanosporofaciens*, whose biosynthesis is catalyzed by four enzymes, CotB1 through CotB4 [15] (Fig 9). CotB1 synthesizes geranylgeranyl diphosphate (GGDP), while CotB2, the main terpene synthase, catalyzes the cyclization and hydroxylation of GGDP to produce cyclooctat-9-en-7-ol (Fig 9B). Subsequently, CotB3 and CotB4, both P450 hydroxylases, introduce hydroxyl groups at the C5 and C18 positions, respectively, to yield cyclooctatin (Fig 9B).

**Fig 9.**
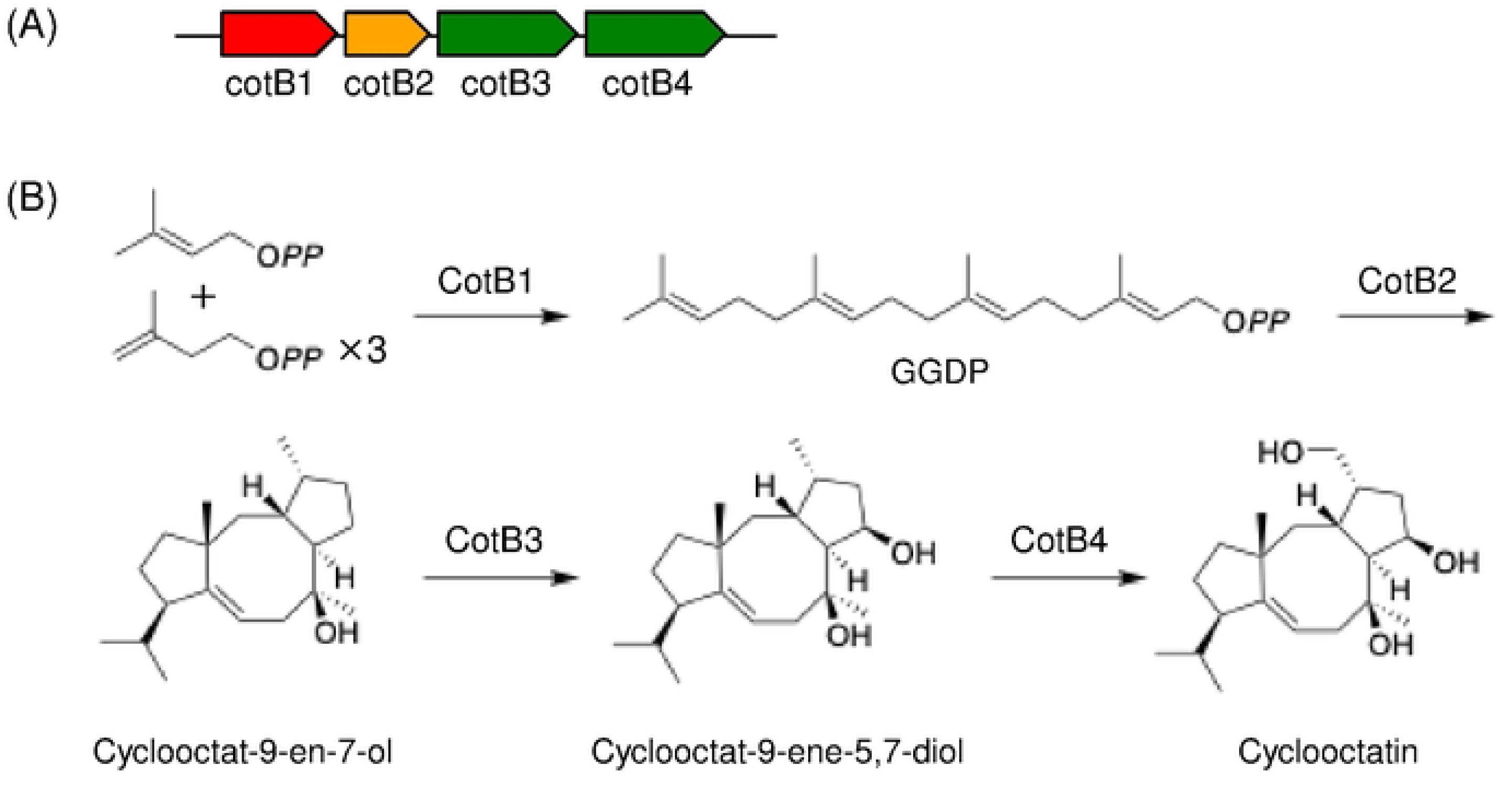
Cyclooctatin biosynthetic gene cluster and biosynthetic mechanism. (A) Gene cluster responsible for cyclooctatin biosynthesis, comprising four genes, *cotB1* to *cotB4*. (B) Mechanism of cyclooctatin biosynthesis [15].

When introducing a masked domain after the CotB4 gene in the cyclooctatin BGC, the genome-trained model (Model IV) predicted functional domains such as FAD_binding_3, Bac_luciferase, and Glyoxalase, which were not predicted by the BGC-trained model (Model I) (Table 3). These newly predicted domains represent candidate functions for modifying cyclooctatin or its intermediates.

**Table 3.**
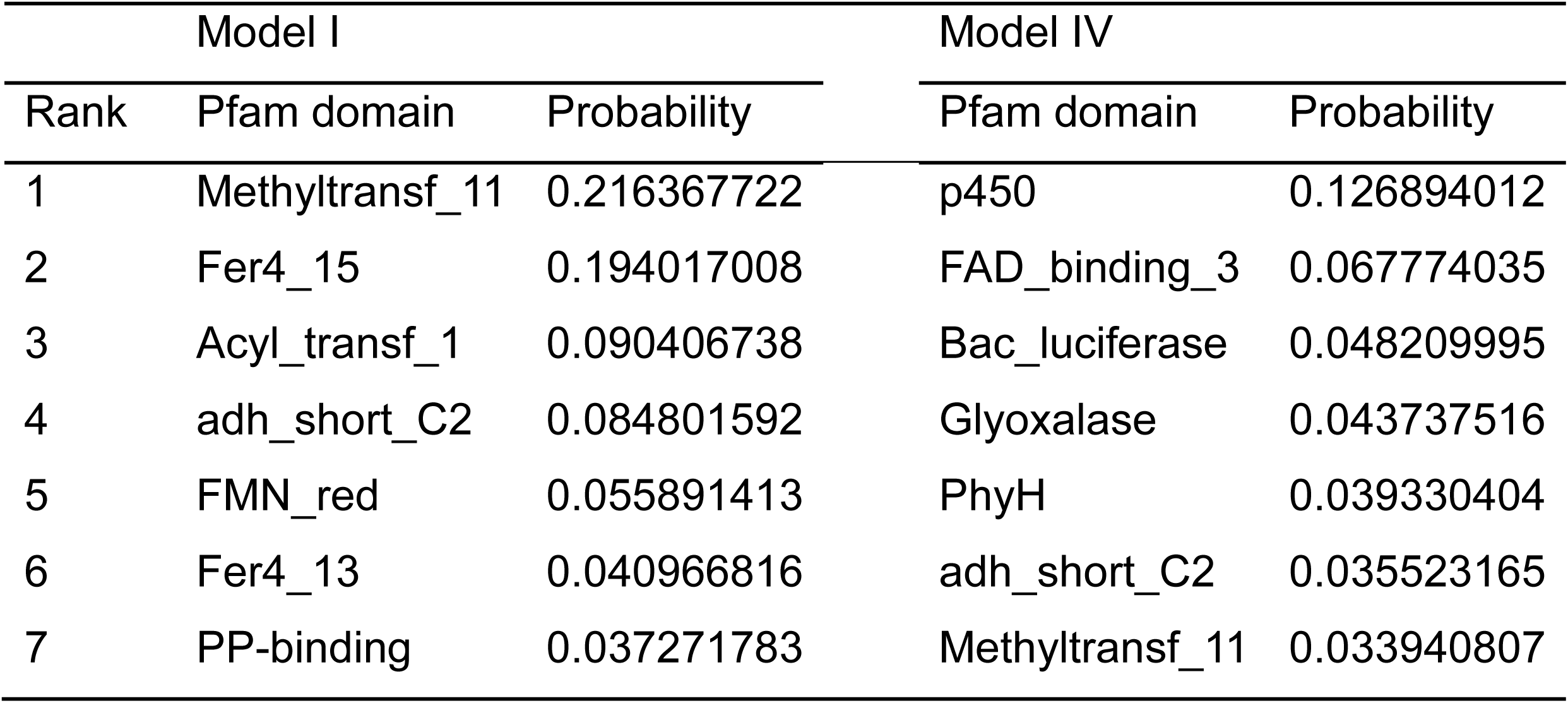
Comparison of top-7 predicted domains for an additional domain to the cyclooctatin biosynthetic gene cluster by Model I and Model IV.

| Model I |  |  | Model IV |  |
| --- | --- | --- | --- | --- |
| Rank | Pfam domain | Probability | Pfam domain | Probability |
| 1 | Methyltransf_11 | 0.216367722 | p450 | 0.126894012 |
| 2 | Fer4_15 | 0.194017008 | FAD_binding_3 | 0.067774035 |
| 3 | Acyl_transf_1 | 0.090406738 | Bac_luciferase | 0.048209995 |
| 4 | adh_short_C2 | 0.084801592 | Glyoxalase | 0.043737516 |
| 5 | FMN_red | 0.055891413 | PhyH | 0.039330404 |
| 6 | Fer4_13 | 0.040966816 | adh_short_C2 | 0.035523165 |
| 7 | PP-binding | 0.037271783 | Methyltransf_11 | 0.033940807 |

### Experimental Validation of Model Prediction

To experimentally validate these model-generated domain suggestions, we need to obtain appropriate amino acid sequences of proteins containing suggested domains, because currently our model solely depends on domain sequences. For this purpose, we used the EFI-GNT web resource [16,17] to search BGCs containing both a CotB1 homolog and a gene encoding domains uniquely predicted by Model IV. We identified such candidate clusters containing the FAD_binding_3, Bac_luciferase, or Glyoxalase domains; however, in the latter two cases, the predicted domain-containing genes were separated by at least two intervening genes from the CotB1 homolog. To prioritize contiguous gene arrangement, we selected a BGC from *Streptomyces* sp. ISL86, which possesses a CotB1 homolog (E-value: 5e-119) and a gene encoding the predicted domain, separated by one intervening gene from the CotB1 homolog.

Based on the amino acid sequence translated from the gene encoding the FAD_binding_3 domain, we constructed a plasmid harboring this gene and co-expressed it in *S. albus* G153 with a plasmid harboring CotB1 and CotB2, CotB1 through CotB3, or CotB1 through CotB4 (Fig 10A). Subsequent LC-MS analysis revealed trace amounts of unknown metabolites specifically produced by the *S. albus* G153 transformant co-expressing FAD_binding_3 with CotB1 and CotB2, CotB1 through CotB3, or CotB1 through CotB4 (Fig 10B, Fig. S2). High-resolution MS analysis deduced that one of the unknown metabolites shares the same molecular formula as cyclooctat-9-ene-5,7-diol (Figure 10C). Nevertheless, its retention time under the chromatographic conditions was clearly different, suggesting that it is a distinct structural isomer of cyclooctat-9-ene-5,7-diol.

**Fig 10.**
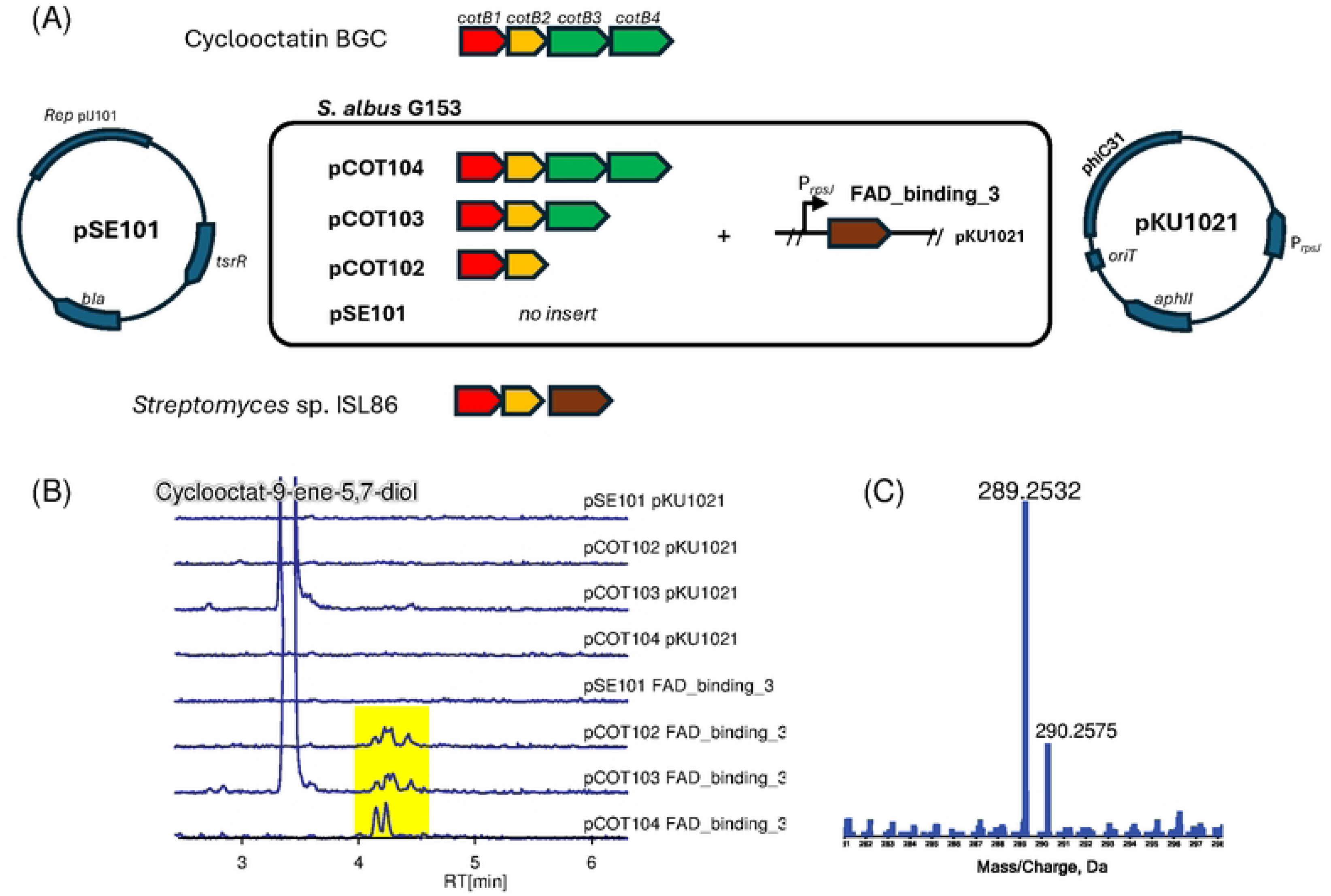
Functional analysis of the domain predicted by the model. (A) A plasmid encoding the gene with the predicted FAD_binding_3 domain from *Streptomyces* sp. ISL86 was constructed and co-expressed with plasmids harboring *cotB1* to *cotB4* in *S. albus* G153. (B) Extracted ion chromatograms for *m/z* 289.252 in the liquid chromatography–mass spectrometry analysis. The culture extracts were analyzed from the *S. albus* transformants harboring pCOT102/103/104 with and without the FAD_binding gene. Cyclooctat-9-ene-5,7-diol (*m/z* 289.2526 [M−H_2_O+H]^+^), highlighted in yellow, was observed to elute at 3.3 min in the the extracts from *S. albus* pCOT103 pKU1021 and *S albus* pCOT103 FAD_binding_3. Although detected in only trace amounts, unknown metabolites were observed to elute at around 4.2 minutes in *S. albus* pCOT102 FAD_binding_3, *S. albus* pCOT103 FAD_binding_3, and *S. albus* pCOT104 FAD_binding_3. (C) High resolution MS spectrum of one of the unknown metabolites in the extract from *S. albus* pCOT102 FAD_binding_3.

This finding validates the model’s ability to propose novel, biologically plausible domain combinations that can be functionally expressed in a heterologous host.

## Discussion

This study demonstrates that our transformer-based model can effectively learn and predict BGC patterns by treating functional domains as linguistic units. This approach provides both conceptual insights into BGC organization and practical tools for natural product discovery. Notably, divergent prediction patterns between the BGC-trained and genome-trained models suggest that broader training contexts may reveal alternative or previously unexplored biosynthetic trajectories, as exemplified in the analysis of cyclooctatin BGC.

We experimentally validated a model-predicted domain combination through heterologous co-expression of the FAD_binding_3 gene with cyclooctatin biosynthetic genes. The FAD_binding_3 domain is found in various enzymes, typically monooxygenases and hydroxylases, where it binds and utilizes flavin adenine dinucleotide as a redox cofactor. Although detected in trace amounts, a putative isomer of cyclooctat-9-ene-5,7-diol was specifically observed in transformants co-expressing CotB1-B2, CotB1-B3, or CotB1-B4 with the FAD_binding_3 protein (Fig 10B, Fig. S2). This observation aligns with the model’s prediction that such domain combination could lead to the production of unknown metabolites. While structural elucidation of the unknown isomer remains necessary, this form of metabolomic validation offers critical feedback for iterative model refinement.

Compared to existing tools such as antiSMASH, which rely on known BGC patterns, our model learns from both BGC and non-BGC genomic regions and can propose novel domain combinations. The use of a transformer architecture enables the capture of long-distance dependencies among domains, which is crucial for understanding cluster organization. These strengths are reflected in the models’ high prediction accuracy and its ability to propose biologically plausible, previously uncharacterized domain assemblies.

Nonetheless, several limitations remain. First, although the current model effectively predicts domain-level organization, it does not generate amino acid sequences optimized for protein-protein interactions or catalytic compatibility. This is because the model is based exclusively on genomic co-occurrence patterns, without incorporating nucleotide or amino acid sequence information. We plan to address this issue by integrating protein sequence design tools such as ProteinMPNN [18] or ProGen [19], as well as nucleic acid generation tools such as Evo [9]. Second, the model cannot autonomously generate candidate artificial BGCs; instead, candidate domains must be selected by comparing predictions across models trained on different datasets. This limitation could be overcome by incorporating additional computational modules, including rule-based heuristics for candidate selection and neural network-based components for domain sequence generation and translation.

The model’s ability to predict biologically plausible domain combinations can prioritize BGCs for experimental validation. Differences in predictions between BGC- and genome-trained models may serve as indicators of evolutionarily atypical clusters with potential for novel bioactivity. Future developments should include integrating transcriptomic data to capture co-expression patterns, using protein structure predictions to assess domain compatibility, and incorporating phylogenetic and metabolic network data to improve biological plausibility. These enhancements can be achieved through a pre-training and fine-tuning framework that balances computational efficiency with multimodal data integration.

In conclusion, this study demonstrates the potential of applying natural language modeling techniques to understand and design BGCs. While the current framework focuses on domain-level prediction, future integration of sequence design, gene expression patterns, and structural constraints will enable generation of more evolutionarily informed and functionally robust BGCs. This transformer-based strategy provides a scalable platform for both theoretical studies of BGC evolution and practical applications in the discovery and engineering of natural products.

## Methods

### Data Collection and Preprocessing

#### Genomic Data

Microbial genome data was downloaded from the National Center for Biotechnology Information. For bacterial genomes, we selected 12,186 complete genomes annotated in RefSeq. For fungal genomes, 2,670 assemblies were collected using GenBank submitter annotations. For each genome, both gene location files (GFF format) and protein amino acid sequences (protein-FASTA format) were retrieved.

#### BGC Data

We obtained 239,021 predicted bacterial BGC data (GenBank format) from the antiSMASH database. Protein sequences of genes located within BGC regions were extracted from each file. Each BGC was treated as a linearized sequence of functional domains for model input.

#### Pfam Domain Identification

Pfam domains were annotated using HMMer version 3.3.1 [20] with the Pfam-A.hmm database (Pfam v35.0) [21]. The E-value threshold was set to 1e-10. Domain annotations were integrated with GFF files to determine domain coordinates on genomes. In cases where domain annotations overlapped in a region, the domain with the better alignment score was retained. A special token [SEP] was inserted between genes to preserve gene boundary information in domain sequences.

### Construction of Tokenization System

Pfam domain names were extracted from the Pfam-A.hmm database. A tokenizer containing 19,523 unique domain tokens was constructed using the PreTrainedTokenizerFast module from the HuggingFace Transformers library (https://huggingface.co/docs/transformers/index). Each domain was treated as a discrete token, allowing BGCs and genomic sequences to be represented as tokenized sequences for language model input.

### Model Architecture and Training

#### RoBERTa Model

We used the RoBERTa architecture implemented in the Hugging Face Transformers library. Model hyperparameters were configured as follows:

- Number of hidden layers: 8
- Number of attention heads: 16
- Hidden layer dimension: 1,024
- Maximum input sequence length: 512 tokens
- Total number of trainable parameters: 105,781,515

Training was conducted using the Masked Language Modeling (MLM) tasks. For each input sequence, 15% of tokens were randomly selected for masking, and among them, 80% were replaced with [MASK] tokens, 10% with random tokens, and the remaining 10% were kept unchanged. The model learned to predict the original tokens based on surrounding context.

#### Training Datasets

Four models were trained on datasets with increasing taxonomic scopes:

- Model I: bacterial BGCs
- Model II: Actinomycetes genomes
- Model III: bacterial genomes
- Model IV: bacterial plus fungal genomes

For each dataset, 80% of samples were randomly assigned for training and 20% reserved for testing. All models shared the same architecture and training procedure.

### Evaluation Methods

#### BGC Classification Task

To assess model performance in compound class prediction, a token indicating the BGC class (*e.g.*, T1PKS, T2PKS, NRPS, terpene) according to antiSMASH classification was appended to the beginning of each domain sequence from a dataset including 22,258 Actinomycetes BGCs retrieved from the antiSMASH database. During training, this label token was masked as part of the MLM task.

The model performance was evaluated using predictions for 5,600 Actinomycetes BGCs retrieved from the antiSMASH database, which were not used for the training.

#### Visualization of Feature Representations

To visualize learned feature representations, we performed t-SNE dimensionality reduction using embeddings from Model IV:

1. The domain sequence of each of the 2,492 MIBiG BGCs was input into Model IV.
2. For each domain, a 1,024-dimensional output feature vector from the model was extracted.
3. The domain vectors for each BGC were averaged to obtain a single feature representation per BGC.
4. Feature vectors were compressed to two dimensions using the t-SNE algorithm (scikit-learn implementation, perplexity=30, random_state=42).
5. BGCs were visualized in 2D space, colored according to their compound class annotations.

### Experimental Validation

#### General

Biochemicals and enzymes for genetic manipulation were purchased from NEB (Tokyo, Japan). Oligonucleotide primers used for genetic manipulation were purchased from Fasmac Co., Ltd. (Kanagawa, Japan). HR-ESI-MS spectra were collected using an SCIEX X500R QTOF system equipped with a UPLC Nexera system (Shimadzu, Kyoto, Japan).

#### Strain Construction

The gene encoding the FAD_binding_3 domain was PCR-amplified using the primers FAD3_fw (GCC<u>AAGCTT</u>TCAGGAGCCGGC) and FAD3_rv(TCC<u>TCTAGA</u>ATGCAGCAGCGCCC) from the synthetic gene (synthesised by Twist Bioscience, codon-optimized for expression in *Streptomyces coelicolor* A3(2), Table S2). The PCR-amplified gene was digested with XbaI and HindIII and cloned into the pKU1021 vector to form pKU1021_FAD_binding_3. The resultant vector was then used for the transformation of *Streptomyces albus* G153 with polyethylene glycol-mediated protoplast transformation. Protoplasts were prepared using the standard protocol. The resultant strain was transformed using pSE101, pCOT102, pCOT103 or pCOT104 with the same method, which were previously constructed [15].

#### Metabolic analysis

Each heterologous expressing strain was inoculated into 10 mL TSB and incubated with shaking (300 rpm) at 30°C for 2 d. A total of 2 mL of the preculture was inoculated into 100 mL of TSB medium and incubated continuously with shaking (180 rpm) at 27°C for 3 d. After fermentation, two times volume of acetone was added to the culture broth. After the extraction, the acetone was evaporated, and the remaining aqueous fraction was extracted with ethyl acetate. The ethyl acetate fraction was recovered and evaporated in vacuo. The remaining residue was dissolved in methanol and analyzed using HR-ESI-LC-MS equipped with a CAPCELL PAK C18 IF column (2.0 φ x 50 mm; Shiseido, Tokyo, Japan). LC conditions were as follows: mobile phase A, water + 0.1% formate; mobile phase B, acetonitrile + 0.1% formate; 10%–90% B over 5 min, 90% B for 2.5 min and then 10% A for 2.5 min, at a flow rate of 0.4 mL/min.

## Acknowledgments

We thank Dr Totai Mitsuyama at National Institute of Advanced Industrial Science and Technology and Dr Yuki Kanai at University of Tokyo for valuable discussion.

## Author Contributions

Tomoki Kawano: Methodology, Data curation, Software, Validation, Investigation, Visualization, Writing – Original Draft.

Taro Shiraishi: Methodology, Validation, Investigation, Writing – Original Draft, Writing - Review & Editing.

Tomohisa Kuzuyama: Project administration, Supervision, Funding acquisition, Writing – Review & Editing.

Maiko Umemura: Conceptualization, Methodology, Supervision, Funding acquisition, Writing – Review & Editing.

## Supporting information captions

Table S1. Domain prediction performance across four training datasets using Actinomycetota 1,042 biosynthetic gene clusters (BGCs) in the MIBiG database.

Table S2. Nucleotide and amino acid sequences of the synthetic gene encoding the FAD_binding_3 domain used in this study.

Fig S1. Training and validation loss curves of RoBERTa and GPT-1 models on Dataset II.

Fig S2. Extracted ion chromatograms for cyclooctatin and its intermediates from Streptomyces albus transformants harboring pCOT102/103/104, with or without the FAD_binding_3 gene.

## Data availability statement

All genomic data, model construction codes, statistical analysis code, and trained models are available on Zenodo at link http://doi.org/10.5281/zenodo.15544024.

## Notes

### Competing Interest Statement

The authors have declared no competing interest.

## References

1. Kanai Y, Tsuru S, Furusawa C. Experimental demonstration of operon formation catalyzed by insertion sequence. Nucleic Acids Res. 2022;50(3):1673–1686. doi:10.1093/nar/gkac004.

2. Schmidt EW, Nelson JT, Rasko DA, Sudek S, Eisen JA, Haygood MG, et al. Patellamide A and C biosynthesis by a microcin-like pathway in Prochloron didemni, the cyanobacterial symbiont of Lissoclinum patella. Proc Natl Acad Sci U S A. 2005;102(20):7315–7320. doi:10.1073/pnas.0501424102.

3. Kenshole E, Herisse M, Michael M, Pidot SJ. Natural product discovery through microbial genome mining. Curr Opin Chem Biol. 2021;60:47–54. doi:10.1016/j.cbpa.2020.07.010.

4. Blin K, Shaw S, Augustijn HE, Reitz ZL, Biermann F, Alanjary M, et al. antiSMASH 7.0: new and improved predictions for detection, regulation, chemical structures and visualisation. Nucleic Acids Res. 2023;51(W1):W46–W50. doi:10.1093/nar/gkad344.

5. Hannigan GD, Prihoda D, Palicka A, Soukup J, Klempir O, Rampula L, et al. A deep learning genome-mining strategy for biosynthetic gene cluster prediction. Nucleic Acids Res. 2019;47(18):e110. doi:10.1093/nar/gkz654.

6. Jumper J, Evans R, Pritzel A, Green T, Figurnov M, Ronneberger O, et al. Highly accurate protein structure prediction with AlphaFold. Nature. 2021;596:583–589. doi:10.1038/s41586-021-03819-2.

7. Rives A, Meier J, Sercu T, Goyal S, Lin Z, Liu J, et al. Biological structure and function emerge from scaling unsupervised learning to 250 million protein sequences. Proc Natl Acad Sci U S A. 2021;118(15):e2016239118. doi:10.1073/pnas.2016239118.

8. Zhang S, Fan R, Liu Y, Chen S, Liu Q, Zeng W. Applications of transformer-based language models in bioinformatics: a survey. Bioinformatics Adv. 2023;3(1):vbad001. doi:10.1093/bioadv/vbad001.

9. Nguyen E, Chen W, Li Y, Wang Y, Zhang X, Zhao H, et al. Sequence modeling and design from molecular to genome scale with Evo. Science. 2024;386:eado9336. doi:10.1126/science.ado9336.

10. Liu Y, Ott M, Goyal N, Du J, Joshi M, Chen D, et al. RoBERTa: a robustly optimized BERT pretraining approach. arXiv. 2019;arXiv:1907.11692.

11. Devlin J, Chang MW, Lee K, Toutanova K. BERT: pre-training of deep bidirectional transformers for language understanding. arXiv. 2018;arXiv:1810.04805.

12. Vaswani A, Shazeer N, Parmar N, Uszkoreit J, Jones L, Gomez AN, et al. Attention is all you need. arXiv. 2017;arXiv:1706.03762.

13. Terlouw BR, Blin K, Navarro-Muñoz JC, Avalon NE, Chevrette MG, Chavali AK, et al. MIBiG 3.0: a community-driven effort to annotate experimentally validated biosynthetic gene clusters. Nucleic Acids Res. 2023;51(D1):D603–D610. doi:10.1093/nar/gkac1049.

14. van der Maaten L, Hinton G. Visualizing data using t-SNE. J Mach Learn Res. 2008;9:2579–2605.

15. Kim SY, Zhao P, Igarashi M, Sawa R, Tomita T, Nishiyama M, et al. Cloning and heterologous expression of the cyclooctatin biosynthetic gene cluster afford a diterpene cyclase and two P450 hydroxylases. Chem Biol. 2009;16(7):736–743. doi:10.1016/j.chembiol.2009.06.007.

16. Zallot R, Oberg N, Gerlt JA. The EFI web resource for genomic enzymology tools: leveraging protein, genome, and metagenome databases to discover novel enzymes and metabolic pathways. Biochemistry. 2019;58(41):4169–4182. doi:10.1021/acs.biochem.9b00735.

17. Oberg N, Zallot R, Gerlt JA. EFI-EST, EFI-GNT, and EFI-CGFP: Enzyme Function Initiative (EFI) web resource for genomic enzymology tools. J Mol Biol. 2023;435(2):168018. doi:10.1016/j.jmb.2023.168018.

18. Dauparas J, Anishchenko I, Bennett N, Bai H, Ragotte RJ, Milles LF, et al. Robust deep learning-based protein sequence design using ProteinMPNN. Science. 2022;378(6615):49–56. doi:10.1126/science.abn2100.

19. Madani A, McCann B, Naik N, Keskar NS, Anand N, Eguchi RR, et al. ProGen: language modeling for protein generation. Nat Mach Intell. 2023;5:316–329. doi:10.1038/s42256-023-00650-9.

20. Finn RD, Clements J, Eddy SR. HMMER web server: interactive sequence similarity searching. Nucleic Acids Res. 2011;39(suppl_2):W29–W37. doi:10.1093/nar/gkr367.

21. Finn RD, Mistry J, Tate J, Coggill P, Heger A, Pollington JE, et al. The Pfam protein families database. Nucleic Acids Res. 2010;38(Database issue):D211–D222. doi:10.1093/nar/gkp985.

